# Development of a *Plasmodium vivax* biobank for functional *ex vivo* assays

**DOI:** 10.1101/2023.03.17.533128

**Authors:** Rashmi Dash, Kristen M. Skillman, Ligia Pereira, Anjali Mascarenhas, Sheena Dass, Jayashri Walke, Anvily Almeida, Mezia Fernandes, Edwin Gomes, John White, Laura Chery-Karschney, Anar Khandeparkar, Pradipsinh K. Rathod, Manoj T. Duraisingh, Usheer Kanjee

## Abstract

**Background:** *Plasmodium vivax* is the second most prevalent cause of malaria yet remains challenging to study due to the lack of a continuous *in vitro* culture system, highlighting the need to establish a biobank of clinical isolates with multiple freezes per sample for use in functional assays. Different methods for cryopreserving parasite isolates were compared and subsequently the most promising one was validated. Enrichment of early- and late-stage parasites and parasite maturation were quantified to facilitate assay planning.

**Methods:** In order to compare cryopreservation protocols, nine clinical *P. vivax* isolates were frozen with four glycerolyte-based mixtures. Parasite recovery post thaw, post KCl-Percoll enrichment and in short-term *in vitro* culture was measured via slide microscopy. Enrichment of late-stage parasites by magnetic activated cell sorting (MACS) was measured. Short and long-term storage of parasites at either -80°C or liquid nitrogen were also compared.

**Results:** Of the four cryopreservation mixtures, one mixture (glycerolyte:serum:RBC at a 2.5:1.5:1 ratio) resulted in improved parasite recovery and statistically significant (P<0.05) enhancement in parasite survival in short-term *in vitro* culture. A parasite biobank was subsequently generated using this protocol resulting in a collection with 106 clinical isolates, each with 8 vials. The quality of the biobank was validated by measuring several factors from 47 thaws: the average reduction in parasitemia post-thaw (25.3%); the average fold enrichment post KCl-Percoll (6.65-fold); and the average percent recovery of parasites (22.0%, measured from 30 isolates). During short-term *in vitro* culture, robust maturation of ring stage parasites to later stages (>20% trophozoites, schizonts and gametocytes) was observed in 60.0% of isolates by 48 hours. Enrichment of mature parasite stages via MACS showed good reproducibility, with an average 30.0% post-MACS parasitemia and an average 5.30 × 10^5^ parasites/vial. Finally, the effect of storage temperature was tested, and no large impacts from short-term (7 day) or long term (7 – 10 year) storage at -80°C on parasite recovery, enrichment or viability was observed.

**Conclusions:** Here, an optimized freezing method for *P. vivax* clinical isolates is demonstrated as a template for the generation and validation of a parasite biobank for use in functional assays.

## BACKGROUND

*Plasmodium vivax* is the most geographically widespread malaria parasite species and is responsible for the second highest burden of malaria globally. While the total incidence of *P*. vivax has decreased globally from 20.5 million cases in 2000 to 4.9 million cases in 2021 [1], significant obstacles remain in efforts to eradicate this pathogen, including the lack of a *P. vivax* vaccine and increasing resistance to antimalarial drugs.

Research into the fundamental biology of *P. vivax* has been severely limited due to the lack of a continuous *in vitro* culture system [2-4]. This lack of continuous culture necessitates the use of clinical *P. vivax* isolates for experimental investigations, and limited access to these isolates has hampered progress in advancing knowledge about this parasite. Recently, short-term *in vitro* culture approaches for *P. vivax* have been established (including in some cases the use of cryopreserved isolates) [3, 5, 6], making it possible to perform single cycle drug resistance assays [3, 7] as well as invasion assays to assess host reticulocyte tropism [8], the effects of invasion inhibitory antibodies [9-16] and host receptor mutants [17, 18]. There is increasing recognition of the importance of biobanking infectious disease samples [19-21] including *P. falciparum* [22-24]. The generation of a biobank of cryopreserved *P. vivax* isolates that can be reliably thawed for downstream experimentation (functional assays, gene expression assays, genomics) would be a critical resource for further advancements in the field[4].

Cryopreservation of blood stage malaria parasites has a long history with early studies relying on rapid freezing to dry-ice or liquid nitrogen temperatures and rapid thawing of RBCs [25, 26] (reviewed in [27]). These early experiments were performed without cryoprotectants and resulted in low recovery of parasites and high levels of RBC lysis. Most modern methods use glycerol-based cryoprotectant solutions which improved recovery of RBCs [28-31], while a minority have used dimethyl sulfoxide (DMSO) [30, 31] (**Supplementary Table 1**). Several cryopreservation methods for *P. vivax* have been reported [5, 7, 32-36], the majority based on adding various ratios of buffered 57% glycerol/lactate solution (Glycerolyte 57) to packed RBCs. To date however, there has not been a systematic comparison of the effectiveness of these different cryopreservation methods for *P. vivax*, largely due to limited research material from clinical isolates.

Here, a continuous source of patient *P. vivax* isolates from the Goa Medical College and Hospital in Goa, India was utilized to compare the effectiveness of four different cryopreservation approaches. An experimental biobank of cryopreserved *P. vivax* isolates was then established, and these samples were used to validate the efficiency of parasite recovery post thaw, the enrichment of early stage parasites via KCl-Percoll density sedimentation [3, 37], the maturation of parasites in short-term *in vitro* culture and the enrichment of mature stage parasites post MACS [38]. Furthermore, the effects of temperature cycling between -80°C (dry ice/freezer) to -196°C (liquid nitrogen) were tested to model the temperature changes that occur during transportation of frozen samples for short-term and long-term storage. These results will inform future cryopreservation and transportation of *P. vivax* samples and aid in establishment of cryopreserved biobanks for future experimental endeavors with this understudied parasite.

## METHODS

### Ethics approval

The human subject protocol and consent forms for enrolling *Plasmodium*-infected patients in this study at Goa Medical College and Hospital (GMC) were reviewed and approved by the Institutional Review Boards of the Division of Microbiology and Infectious Diseases (DMID) at the U.S. National Institute of Allergy and Infectious Diseases (approval DMID 11-0074), the University of Washington (approval 42271/1192), as well as the Institutional Ethics Committee (IEC) at Goa Medical College Hospital, Bambolim, Goa, India.

### Sample collection

The study was conducted at the Goa Medical College Hospital, at the Malaria Evolution in South Asia-International Centers for Excellence in Malaria Research laboratory. Venous blood was collected in 6 mL vacutainer tubes (acid dextrose solution anticoagulant; BD India) from all the smear positive patients for *Plasmodium vivax* mono infection with signed informed consent between the age of 12 months to 65 years between 2013 to 2019. Pregnant (self-reported) and anemic patients were excluded from enrollment.

### Sample processing & cryopreservation

Clinical samples that tested positive for *P. vivax* by rapid diagnostic test (FalciVax, Zephyr Biomedicals, Goa, India) were collected and smears were prepared and stained using Field Stains A and B (Himedia). Samples with parasitemia >0.01% were included for further analysis. The blood was centrifuged at 800 *g* for 5 minutes, and serum from each isolate was separated and stored at -80°C for related studies. The pelleted blood was washed using phosphate buffered saline (Thermo Scientific) supplemented with 0.5% (w/v) bovine serum albumin (Thermo Scientific) to separate any leftover serum components. Prior to cryopreservation, the pelleted blood was separated from leukocytes, by running the sample through CF-11 column filters prepared in the lab. Smears were prepared post processing with CF-11 to assess parasite stages and to examine the leukocyte removal. De-leukocyted samples were cryopreserved using Glycerolyte 57 (Baxter), added as follows: glycerolyte 57:serum:RBC ratio was 2:0:1 for mixture 1 [7], 2.5:1.5:1 for mixture 2 [35, 39], 1.66:0:1 for mixture 3 [36] and 4.15:1.5:1 for mixture 4. Sample were stored at -80°C for 24 hours prior to long-term storage in liquid nitrogen cryo-tanks at -196°C. The serum used in the study was heat inactivated AB+ pooled plasma (Access Biologicals, USA) collected from a non-malaria endemic region.

### Short term *P. vivax* culture

*Plasmodium vivax* isolates were thawed using a standard protocol [40]. Samples were thawed out rapidly in 37°C water bath followed by dropwise addition of 1/10^th^ volume of 12% (w/v) NaCl and 10 volumes of 1.6% (w/v) NaCl. After washing 2X with IMDM+glutamax media (Thermo Scientific), isolates were enriched using 1.080 g/mL KCL Percoll gradients [3, 37]. Three milliliters of the sample in media were layered on an equal volume of 1.080 g/mL KCl Percoll at room temperature in a 15 mL centrifuge tube. The layered sample was centrifuged at 1200 *g* for 15 minutes with slow acceleration and no brake. The enriched parasites from the Percoll interface were washed in IMDM+glutamax + 0.5% (w/v) BSA and were subsequently cultured in IMDM + 10% (v/v) AB+ heat inactivated serum with 0.5%(v/v) gentamycin per condition in 1 mL volumes in 24-well plates (Nunc). The plates were incubated at 37°C and with mixed gas (5% CO_2_, 10% O_2_ and 95% N_2_ combination) in modular incubator chambers (Billups Rothenberg) for 24-72 hours as needed for the data assessment. Enrichment of parasites via MACS was performed as previously reported [41].

### Microscopy

Slides for the experiments were counted on light microscope using 100X objective with oil immersion with a 1:4 reticle on a Zeiss Primo Star microscope. The slides were counted and accessed for stages using the whole field method [42] across multiple fields (15-20 fields) until 500 red blood cells were counted in the small box of the 1:4 reticle. The slides for the different time point during the experiment were prepared using a cytospin (Shandon/Thermo Scientific) with 100 µL 10% (w/v) bovine serum albumin in PBS. Parasitemia was calculated as:

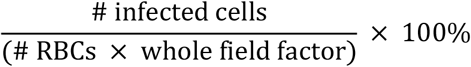

where the whole field factor for the 1:4 reticle was 10.9.

### Data Analysis

All the data for the experiments was analyzed using Microsoft Excel and Graphpad Prism (version 9.8.14). Comparisons between experimental conditions was performed via one-way ANOVA analysis (or where slide data was missing due to unreadable slides via mixed-effect model analysis). The analysis was performed with matching by parasite isolate, with the assumption of a Gaussian distribution of residuals and assumption of sphericity. Tukey’s multiple comparisons test was used to determine statistically significant differences between individual sets of conditions.

## RESULTS

### Comparing cryopreservation mixtures

Among the malaria positive individuals presenting at Goa Medical College and Hospital (GMC), >80% are infected with *P. vivax* [43]. The ability to collect this number of *P. vivax* isolates has made GMC a unique setting to develop a biobank of cryopreserved parasites for future experimentation. From 2012 – 2016, over 700 *P. vivax* isolates were cryopreserved using the method from Kosaisavee *et al*., [7] where a 2:1 ratio of the cryoprotectant glycerolyte:RBCs was used. Other ratios of glycerolyte:RBCs have been reported in the literature for cryopreservation of both *P. falciparum* and *P. vivax* (**Supplementary Table 1**). We decided to test four cryopreservation mixtures, derived from established cryopreservation protocols [7, 34, 36, 39, 44], (**Figure 1A**) that varied in the ratios of glycerolyte:RBCs (with or without the addition of serum) in order to test for any differences with the established 2:1 glycerolyte:RBC ratio.

**Figure 1:**
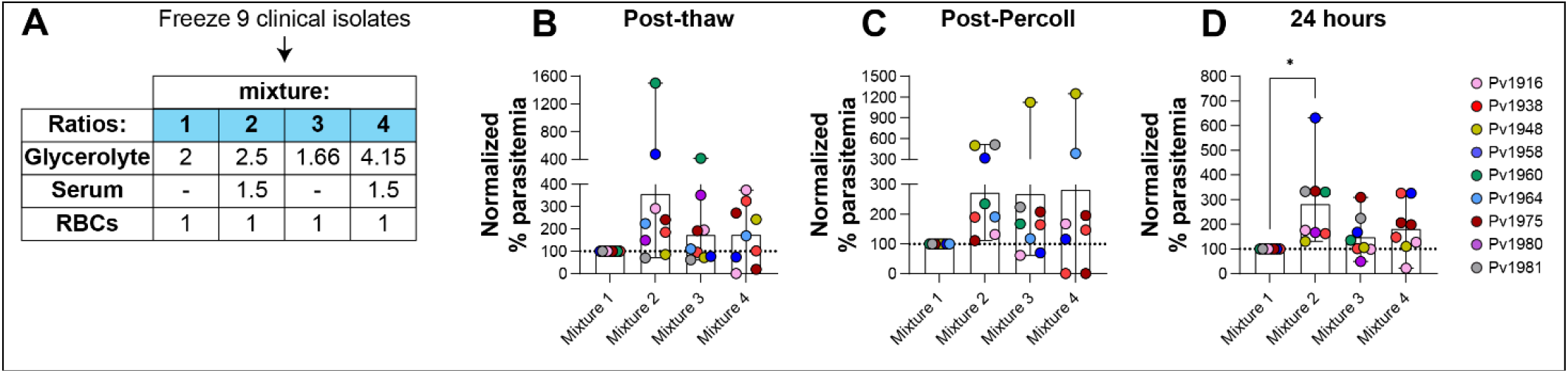
Comparison of four different freezing mixtures. (A) Nine clinical *P*. vivax isolates were cryopreserved using four mixtures with different ratios of RBCs, serum and glycerolyte. Comparison of (B) post-thaw parasitemia, (C) post KCl-Percoll parasitemia and (D) 24-hour parasitemia all normalized to mixture 1 values. * P<0.05 via one-way ANOVA with Dunnett’s multiple comparison test.

Nine *P. vivax* isolates were split into 4 equal portions, and each was cryopreserved using one of the mixtures. After a minimum of 1 week’s storage in liquid nitrogen, these samples were thawed using standard methods [40] and the parasitemia determined via slide microscopy post thaw, post enrichment via KCl-Percoll density gradients [3, 37] and at 24 hours in short-term *ex vivo* culture. Parasitemia values were normalized to the mixture 1 values. Post thaw (**Figure 1B**), an increase in relative parasitemia was observed for mixture 2 but this did not reach statistical significance. Post KCl-Percoll enrichment (**Figure 1C**), mixtures 2, 3 and 4 showed higher relative parasitemia than mixture 1, but again did not reach statistical significance. At 24 hours in short-term culture (**Figure 1D**), a statistically significant increase in relative parasitemia was observed for mixture 2 compared to mixture 1 (P = 0.0123, F (1.781, 12.47) = 6.690 via one-way ANOVA with matched data and Dunnett’s multiple comparisons test). Together these data demonstrated that cryopreservation of *P. vivax* isolates with mixture 2 resulted in improved recovery and maturation of parasites and this cryopreservation formulation was chosen for the establishment of an improved biobank.

### Generation and validation of a biobank

Over one hundred *P. vivax* isolates have been cryopreserved using mixture 2, with an average of 8 vials generated per sample (∼ 300 µL packed RBCs/vial). The initial parasitemias ranged from 0.036% to 1.526% with a median of 0.323% and average of 0.384% (**Figure 2A**). Within the collected samples, the mean parasitemia values were highest for ring stages (early rings (0.212%); late rings (0.071%)) compared to other stages: early trophozoites (0.044%); late trophozoites (0.008%); schizonts (0.005%) and gametocytes (0.040%) (**Figure 2B**).

**Figure 2:**
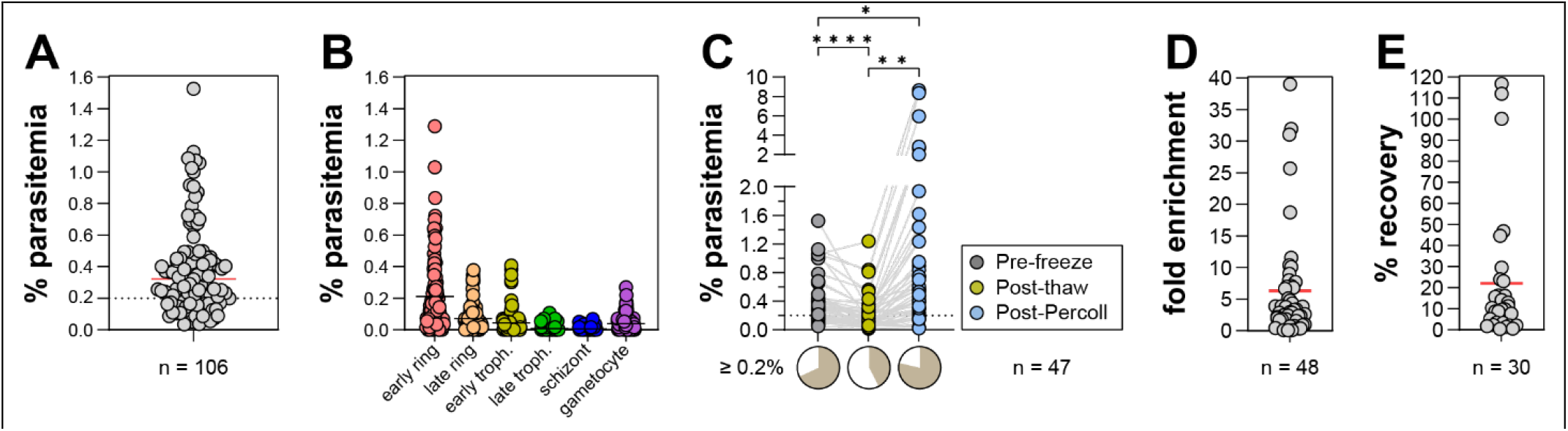
Characterization of biobank and enrichment of rings via KCl-Percoll. (A) Range of pre-freeze parasitemia values for 106 cryopreserved isolates. (B) Distribution of different stages pre-freeze. (C) Comparison of parasitemia values pre-freeze, post-thaw and post-KCl-Percoll for 47 isolates. *P<0.05; **P<0.01; ***P<0.001 via One-way ANOVA with Tukey’s multiple comparisons. Lower pie charts show proportion of samples with parasitemias ≥ 0.2%. (D) Distribution of fold-enrichments post KCl-Percoll. (E) Distribution of the percent recovery of parasites during KCl-Percoll enrichment.

Both drug resistance and invasion assays with *P. vivax* isolates require a sufficient number of parasites for reliable measurements, and as such different strategies have been employed to enrich for parasites either at the ring stage (e.g., for drug resistance assays via KCl-Percoll gradients [3, 7]) or at the schizont stage (e.g., for invasion assays using either Percoll gradients [10] or magnetic enrichment [18, 41, 45]). In a previously reported drug resistance assay with *P. vivax* [3], a minimum of 0.2% parasitemia was necessary for good quality measurements, and we found that 72% of total isolates met that threshold pre-freeze. In order to characterize any changes in parasitemia post-thaw and to measure the effect of KCl-Percoll enrichment, the parasitemia was measured pre-freeze, post-thaw, and post KCl-Percoll enrichment from 47 isolates (**Figure 2C**). A statistically significant difference in percent parasitemia was observed between all three conditions (P = 0.0011 (F (1.034, 47.57) = 11.85) via one-way ANOVA with Tukey’s multiple comparisons). There was an average 74.7% (range 7.21 – 419.36%) recovery of parasitemia between pre-freeze versus post thaw and an average of 6.7-fold (range 0.13 – 39.0-fold) enrichment of parasitemia post KCl-Percoll (**Figure 2D**). Furthermore, the proportion of parasite isolates reaching the 0.2% threshold decreased from 68% pre-freeze to 43% post-thaw but was subsequently increased to 79% post KCl-Percoll (**Figure 2C**, lower pie charts) demonstrating the utility of this method for increasing parasitemia. The percent recovery of parasites during KCl-Percoll enrichments was also estimated by measuring both the percent parasitemia via slide microscopy and the total RBC numbers via hemocytometer counts both post-thaw and post KCl-Percoll (**Figure 2E**). The average recovery was 22% (range 0.43 – 116.8%) suggesting that not all parasites are enriched via this approach.

### Maturation of parasites in short-term *ex vivo* culture

While there is no long-term culture system for *P. vivax*, short-term *ex vivo* maturation of parasites has been reported [3, 6, 7, 10, 41]. In order to establish the ability of biobanked isolates to mature, a total of twenty isolates were selected for validation experiments. The samples were thawed and enriched on KCl-Percoll gradients and the staging and parasitemia were measured post-enrichment and at 24 hours and 48 hours in short-term culture (**Figure 3A**). (During the course of these experiments occasional slides were unreadable/damaged, and these isolates were excluded from analysis at the given timepoint). During the first 24 hours, the average parasitemia remained relatively consistent, with some isolates showing decreases; however, by 48 hours there was an average 64.9% decrease in percent parasitemia across the samples. The maturation of the parasites was quantified by determining the number of isolates with different proportions (<20%, 20–50%, >50%) of either immature parasites (early and late rings) or maturing parasites (trophozoites, schizonts and gametocytes) (**Figure 3B**). Post enrichment, all isolates had at least 20% immature parasites. At 24 hours, 57.9% (11/19) had >20% mature stages, and a subset of 26.3% (5/19) had >50% mature stages. These proportions did not change appreciably as by 48 hours, 60% of isolates (12/20) had >20% mature stages with 30% (6/20) having >50% mature stages.

**Figure 3:**
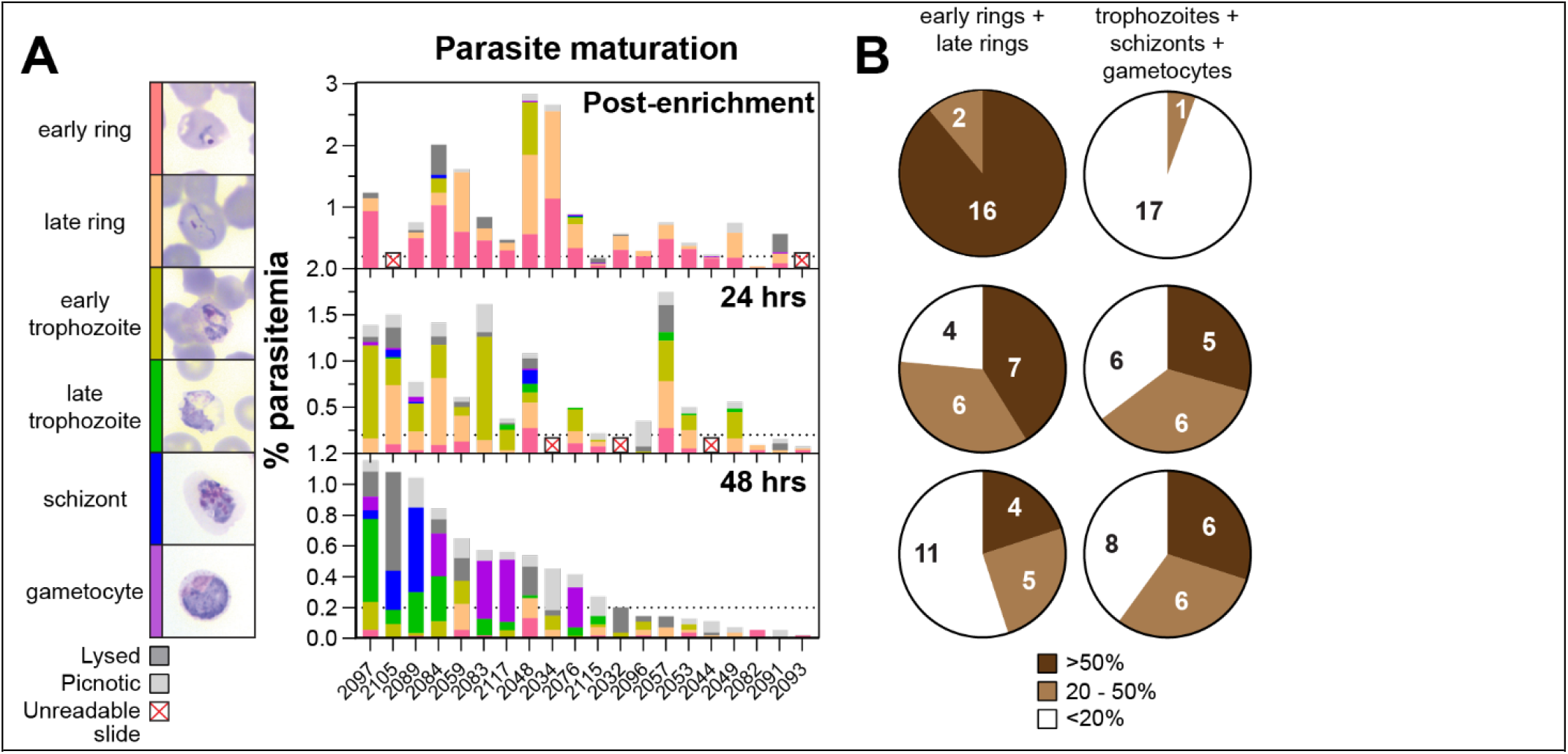
Maturation of parasites in short-term *ex vivo* culture. (A) Staging and parasitemia from 20 biobanked samples post-enrichment, 24-hours, and 48-hours in short-term culture. (B) Pie charts showing proportion of either ring stages (early and late rings) or late-stage (trophozoites, schizonts, gametocytes) parasites across isolates.

### Enrichment of mature stages via MACS

Mature parasites can be readily enriched via magnetic activated cell sorting (MACS) through the paramagnetic properties of hemozoin in the parasite food vacuole [38], and is a common method of purifying parasites for use in invasion assays [18, 41, 45]. The range of parasitemias post MACS enrichment was measured (**Figure 4A**), with an average 29.95% parasitemia (range 0.52% - 80.81%). In addition, the number of parasites per vial was determined (**Figure 4B**), with an average of 5.30 × 10^5^/vial (range 7.68 × 10^4^ – 2.90 × 10^6^/vial). These values would be useful when using parasites for invasion assays in order to determine the number of assay wells that can be usefully set up per isolate.

**Figure 4:**
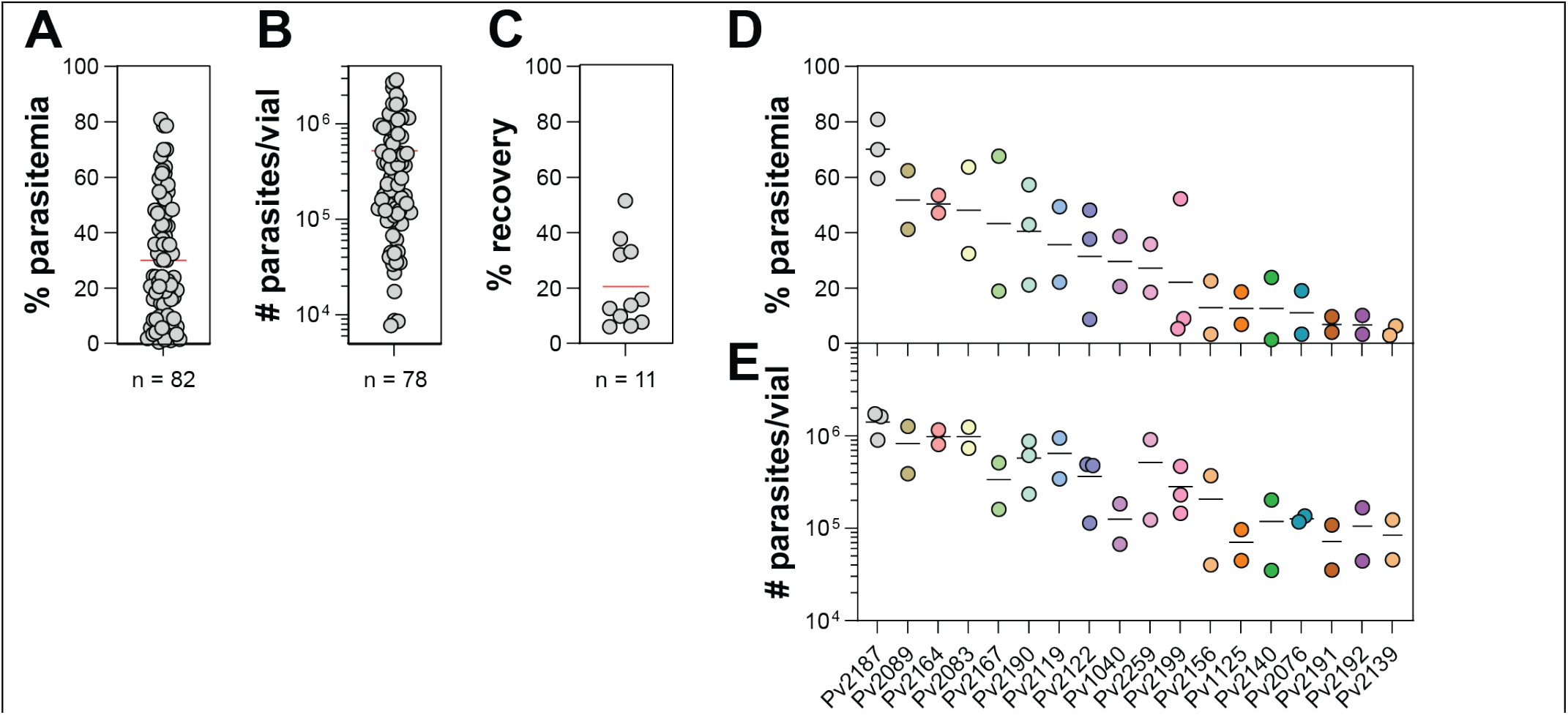
Enrichment of parasites via MACS. (A) Distribution of parasitemias post-MACS enrichment. (B) Calculated number of parasites/vial post-MACS. (C) Calculated % recovery of parasites post-MACS enrichment. Distribution of (D) parasitemia and (E) number of parasites/vial from repeat thaws of biobanked isolates.

The recovery of parasites via MACS enrichment was also estimated by measuring the number of RBCs via hemocytometer and the % parasitemia via slide microscopy pre-MACS and post-MACS. For the 11 samples that were tested, an average of 20.59% recovery was observed (range 5.94% – 51.44%) (**Figure 4C**). The reproducibility of MACS enrichment and parasite recovery was also determined when multiple independent thaws of the same isolate were made (**Figure 4D**,**E**), and in most cases the % parasitemia and the number of parasites per vial varied within less than 2-fold.

### Impact of temperature variation during transport

In addition to the method of cryopreservation, the storage and transport temperature experienced by frozen isolates may impact their viability following thaw. Previous studies with *P. falciparum* have demonstrated reduced viability of cryopreserved parasites stored at -70°C compared to liquid nitrogen temperatures [27, 30]. Therefore, we conducted a temperature variation experiment to mimic transport of samples from the site of collection to other locations for experimental use. Nine samples were cryopreserved using mixture 2, generating four vials, each of which was subjected to four different temperature storage conditions (**Figure 5A**). Vial 1 (V1) was the control condition and was thawed directly from liquid nitrogen within a duration of 5-8 days. Vials V2-V4 (test conditions) were transferred from liquid nitrogen to -80°C for 7 days to mimic shipment conditions on dry ice. Vial V2 was thawed after 7 days of storage at -80°C while vials V3 and V4 were returned to liquid nitrogen for a further 7 days to mimic possible temperature fluctuation at a site where samples were returned to liquid nitrogen for storage. Vial V4 was cycled a second time to -80°C prior to thaw while vial V3 was thawed from liquid nitrogen at 7 days.

**Figure 5:**
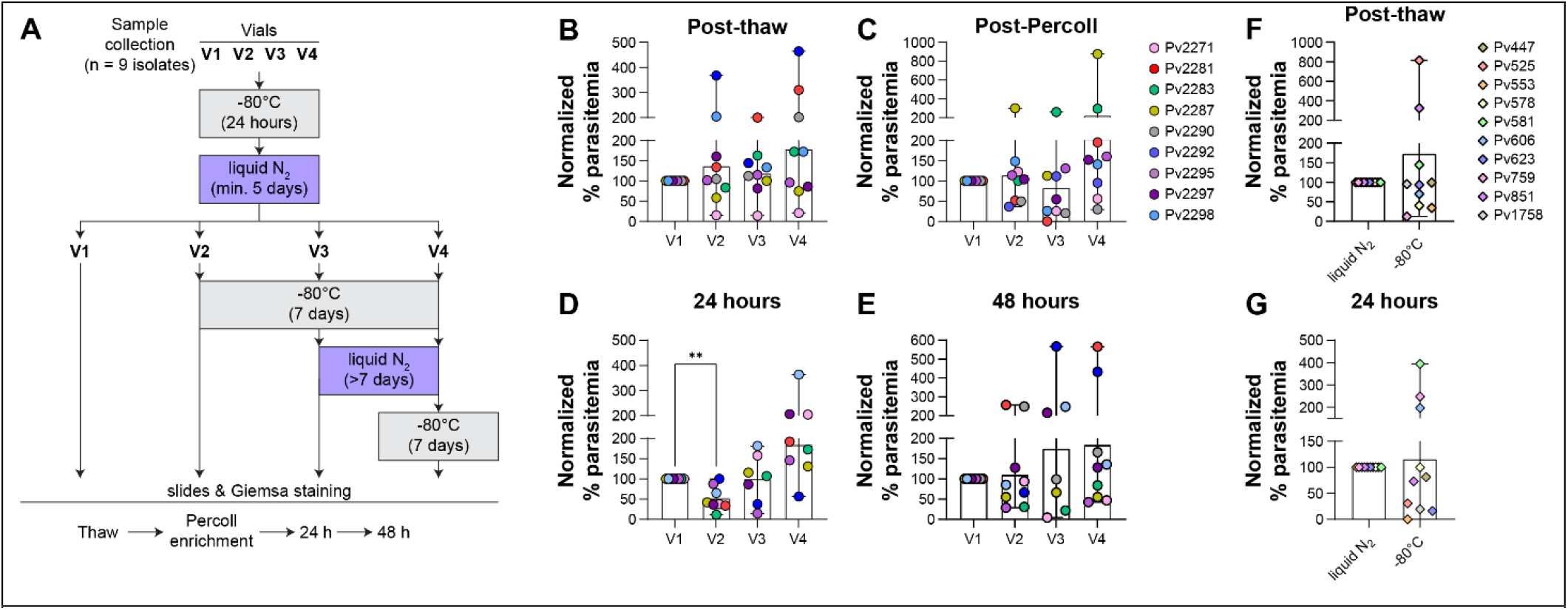
Comparison of freezing and storage conditions. Schematic of parasite freezing and storage conditions tested (V1 – V4). Comparison of samples Post thaw (B), Post-KCl-Percoll (C), 24-hours (D) and 48-hours (E) with parasitemia values normalized to V1. **P<0.01 via mixed effects analysis with Dunnett’s multiple comparisons. Comparison of samples stored long term (7 – 10 years) in either liquid nitrogen or at -80°C: (F) post-thaw and (G) at 24-hours in short-term culture.

Upon thawing, all samples were enriched via KCl-Percoll and short-term cultured with microscopy examination at thaw, enrichment, and following 24 hours and 48 hours *in vitro* growth with data normalized to the V1 values (**Figure 5B-E**). No statistically significant differences were observed between the conditions post-thaw or post-Percoll. However, we did observe a significant reduction in parasitemia for condition V2 at 24 hours (P = 0.010 (F (1.399, 9.330) = 9.024) via multiple effects analysis with Dunnett’s multiple comparisons), although this reduction was not observed at 48 hours. Furthermore, in order to test whether there might be long-term effects of storage of parasites at different temperatures, we compared a set of isolates where vials had been stored at both liquid nitrogen and -80°C temperatures for between 7 – 10 years (**Figure 5F**,**G**) (note that these samples had been cryopreserved using mixture 1). We did not observe any statistically significant difference in parasitemia either post-thaw or at 24 hours in short-term culture, suggesting that either storage temperature is capable of preserving parasite.

## DISCUSSION

The establishment of a biobank of cryopreserved *P. vivax* isolates that can be reliably thawed for downstream use would be a powerful tool to facilitate research with this parasite. The current lack of a culture system for *P. vivax* precludes the type of experiments that can be conducted with other culturable *Plasmodium* species, such as *P. falciparum* [46] and *P. knowlesi* [47-49]. Advances in counting and staging *P. vivax* have been made that would facilitate collection of *P. vivax* for such a resource [3, 42]. Collection of properly preserved and stored isolates, combined with the ability to enrich low parasitemia samples with the use of KCl-Percoll gradients for ring-stage assays (e.g., drug resistance assays) or to enrich mature parasites via Percoll gradients or MACS (e.g., for invasion assays), would facilitate functional assays with this understudied parasite. Not only would more assays be possible, but the generation of a biobank with multiple freezes from the same donor would allow replicate experiments to be conducted, which is generally not possible when only live isolates can be collected. Finally, establishment of a reliable source of cryopreserved isolates extends the duration of experimentation beyond the typical high-transmission peak season when most clinical isolates are collected.

To inform the generation of a cryopreserved biobank, we studied the progression and maturation of asexual blood stages for *P. vivax* following thaws from vials that had been cryopreserved using four different cryopreservation mixtures with different ratios of Glycerolyte 57 to RBCs (with or without the inclusion of serum). Despite the final concentration of glycerol varying (4.12 M (mixture 1), 3.10 M (mixture 2), 3.86 M (mixtures 3 and 4)) (**Supplementary Table 1**), the four mixtures showed broadly similar characteristics in terms of parasitemias post-thaw, post KCl-Percoll enrichment and in short-term *ex vivo* culture. However, mixture 2 was chosen to generate an optimized biobank due to higher recovery and viability of parasites compared to the other mixtures.

Consistent with previous short-term culture optimization experiments [3] we observed a less than 100% recovery of parasites post thaw (**Figure 1C**), as well as the presence of picnotic & lysed parasites during *in vitro* culture (**Figure 3**), suggesting that further improvements in the cryopreservation and thawing methodology may be possible, such as comparing the sorbitol method [50] to see if there is improved recovery of parasites compared to the sodium chloride method currently in use [40].

The use of density gradients to enrich reticulocytes has been reported previously, with focus either on Nycodenz [37], aqueous multiphase systems [51] or Percoll [3, 41, 52] approaches. Here we show the ability of KCl-Percoll density gradients enrich parasites post thaw to parasitemias sufficiently high for drug resistance assays (> 0.2% parasitemia [3]) (**Figure 1C**). The measured recovery of parasites using this method is still relatively low (**Figure 1E**), suggesting that other improvements to this approach may be possible. Future experiments should also directly measure recovery of reticulocytes (both uninfected and infected) as well as the effectiveness of recovery and maturation of sexual stages [53] which may be useful for future transmission studies.

Measurement of the viability and maturation of parasites during short-term *in vitro* culture (**Figure 3**) suggests that at least 60% of isolates in the biobank will be capable of complete maturation (with at least 20% mature stages). We note that this may be an underestimate as schizonts in some isolates may have already matured and egressed by the 48-hour timepoint when slides were made. A more accurate assessment of maturation may require use of egress inhibitors to block parasite egress [54]. The MACS enrichment of mature parasites (trophozoites, schizonts and gametocytes) showed a relatively wide distribution in parasitemias, number of parasites/vial and % recovery post MACS. The less than 100% recovery may be due to the wide distribution of parasite stages observed during short-term ex vivo culture (**Figure 3**). In addition, the enrichment of parasites between independent thaws showed relatively good reproducibility (**Figure 4D**,**E**) allowing for some ability to predict the number of mature parasites expected from an individual thaw which would be key in planning invasion assays [17, 41, 55].

While long-term storage at liquid nitrogen is preferred, little is known about the effect of cycling between liquid nitrogen and dry-ice/-80°C storage temperatures on viability of *Plasmodium* spp. parasites [27]. When parasite isolates are collected from endemic settings specifically, the available options for storage of samples during transport from the collection point to long term storage may be limited. We compared the short-term storage of isolates at different temperature regimes and found no significant difference in parasite viability post-thaw and post enrichment, while we did observe a reduction in parasitemia at 24 hours in short term culture for parasites that had been stored at -80°C (**Figure 5D**). However, comparison of long-term storage of samples in liquid nitrogen and at -80°C did not reveal any statistically significant differences between storage at either temperature, although it is possible that viability may vary from sample to sample.

## CONCLUSIONS

We have compared four different cryopreservation methods for *P. vivax*, and we have found one method to provide improved parasite recovery and maturation in short-term *in vitro* cultures. Using this method, we have established a biobank of clinical isolates with multiple freezes per isolate and we have validated the biobank showing the ability to enrich parasites post thaw with KCl-Percoll gradients, and for at least 60% of parasites to successfully mature in short-term *in vitro* cultures. Finally, we have tested storage of cryopreserved parasites at different temperatures (liquid nitrogen storage and -80°C storage), but at least for the duration of this study, we found no evidence of changes in viability of parasites stored in liquid nitrogen versus - 80°C.

## List of abbreviations

(RBC): Red blood cell
(MACS): Magnetically activated cell sorting
(DMSO): dimethyl sulfoxide
(GMC): Goa Medical College and Hospital

## DECLARATIONS

### Ethics approval

The human subject protocol and consent forms for enrolling *Plasmodium*-infected patients in this study at Goa Medical College and Hospital (GMC), by the Institutional Review Boards of the Division of Microbiology and Infectious Diseases (DMID) at the U.S. National Institute of Allergy and Infectious Diseases (approval DMID 11-0074), the University of Washington (approval 42271/1192), as well as Goa Medical College Hospital, Bambolim, Goa, India.

### Availability of data

The datasets used and/or analyzed during the current study are available from the corresponding author on reasonable request.

### Author’s contributions

RD, LP, AM, JW, AA, MF, UK, SD collected samples and generated data. RD, KMS and UK analyzed data and wrote the manuscript. EG, JW, LCK, PKR, UK, KMS, RD, AK, SD and MDT interpreted and revised the manuscript

## Figures and Tables

**Supplementary Table 1:** Summary of *Plasmodium* spp. cryopreservation methods

| Reference | Year | Species | Freezing Solution | Freezing Solution:RBC ratio | Final concentration of cryoprotectant |
| --- | --- | --- | --- | --- | --- |
| [35] | This study | *P. vivax* | Glycerolyte 57<sup>§</sup> | 1:1 (60% serum; 40% RBC) **(Mixture 2)** | 3.10 M glycerol |
| [36] | This study | *P. vivax* | Glycerolyte 57<sup>§</sup> | 1.66:1 (60% serum; 40% RBC) **(Mixture 4)** | 3.86 M glycerol |
| [36] | 2017 | *P. vivax* | Glycerolyte 57<sup>§</sup> | 1.66:1 | 3.86 M glycerol |
| [35] | 2017 | *P. vivax* | Glycerolyte 57<sup>§</sup> | 1:1 | 3.10 M glycerol |
| [56] | 2015 | *P. falciparum* | Glycerolyte 57<sup>§</sup> | 2:1 | 4.12 M glycerol |
| [48] | 2013 | *P. knowlesi* | Glycerol/sorbitol<sup>‡</sup> | 7:3 | 2.69 M glycerol |
| [34] | 2013 | *P. vivax*; *P. cynomolgi* | Glycerol/mannitol<sup>‡</sup> | 1:1 | 1.92 M glycerol |
| [44] | 2013 | *Plasmodium* spp. | Glycerolyte 57<sup>§</sup> | 1.66:1 **(Mixture 3)** | 3.86 M glycerol |
| [39] | 2013 | *Plasmodium* spp. | Glycerol/sorbitol<sup>‡</sup> | 1:1 (60% serum, 40% RBC) | 1.92 M glycerol |
| [33] | 2012 | *P. vivax* | Glycerolyte 57<sup>§</sup> | 2:1 | 4.12 M glycerol |
| [5] | 2012 | *P. vivax* | Glycerolyte 57<sup>§</sup> | 2:1 | 4.12 M glycerol |
| [7] | 2006 | *P. vivax* | Glycerolyte 57<sup>§</sup> | 2:1 **(Mixture 1)** | 4.12 M glycerol |
| [47] | 2002 | *P. knowlesi* | Glycerol/mannitol<sup>‡</sup> | 1:1 | 1.92 M glycerol |
| [31] | 1989 | *P. falciparum* | Glycerolyte<sup>#</sup> | 4:1 | 4.96 M glycerol |
| [31] | 1989 | *P. falciparum* | 20% (v/v) DMSO | 1:1 | 1.41 M DMSO |
| [32] | 1985 | *P. falciparum*; *P. vivax* | Alsevers solution + 10% (w/v) glycerol | 3:1 | 1.37 M glycerol |
| [29] | 1975 | *P. falciparum* | Glycerolyte<sup>#</sup> | 1.6:1 | 3.82 M glycerol |
| [57] | 1973 | *P. falciparum*, *P. knowlesi* | 15% (v/v) DMSO, 0.85% (w/v) NaCl | 1:1 | 1.06 M DMSO |
| [28] | 1968 | RBCs | Glycerol/mannitol<sup>‡</sup> | 1:1 | 1.92 M glycerol |
| [26] | 1957 | *P. falciparum*; *P. ovale* | None | N/A | N/A |
| [25] | 1939 | *P. knowlesi* | None | N/A | N/A |

<sup>§</sup>Composition of Glycerolyte 57: 6.19 M glycerol; 142.8 mM sodium lactate; 4.0 mM KCl, 3.7 mM NaH<sub>2</sub>PO<sub>4</sub>, 8.7 mM Na<sub>2</sub>HPO<sub>4</sub>, pH 6.8
<sup>#</sup>Composition of Glycerolyte: 6.2 M glycerol, 140 mM sodium lactate, 5 mM KCl, NaH<sub>2</sub>PO<sub>4</sub> to pH 7.4
<sup>‡</sup>Composition of Glycerol/mannitol solution: 28% (w/v) glycerol, 3% (w/v) mannitol, 0.65% (w/v) NaCl
<sup>‡</sup>Composition of Glycerol/sorbitol solution: 28% (w/v) glycerol, 3% (w/v) sorbitol, 0.65% (w/v) NaCl
<sup>¶</sup>Composition of complete RPMI: RPMI 1640 supplemented with 25 mM HEPES, 0.2% (w/v) NaHCO<sub>3</sub>, 10% (v/v) serum

